# Trends in millennial adolescent mental health and health related behaviours over ten years: a population cohort comparison study

**DOI:** 10.1101/407585

**Authors:** Praveetha Patalay, Suzanne H Gage

## Abstract

**Background:** There is evidence that mental health problems are increasing and substance use behaviours are decreasing. This paper aimed to investigate recent trends in mental ill-health and health-related behaviours in two cohorts of UK adolescents in 2005 and 2015.

**Method:** Trends in harmonised mental ill-health (depressive symptoms, self-harm, anti-social behaviours, parent reported difficulties) and health related behaviours (substance use, weight, weight perception, sleep, sexual intercourse) were examined at age 14 in two UK birth cohorts; Avon Longitudinal Study of Parents and Children (ALSPAC, N=5627, born 1991-92) and Millennium Cohort Study (MCS, N=11318, born 2000-02). Prevalences and trend estimates are presented unadjusted and using propensity score matching and entropy balancing.

**Results:** Depressive symptoms (9% to ∼15%) and self-harm (11.8% to ∼14.5%) increased over the 10 years. Parent-reported emotional difficulties, conduct problems, hyperactivity and peer problems were higher in 2015 compared to 2005 (5.7 – 9% to 9 – 18%).Conversely, substance use (tried smoking 9% to 3%; tried alcohol 52% to ∼43%, cannabis 4.6% to ∼4%), sexual activity (2% to ∼1%) and anti-social behaviours (6.2 – 40.1% to 1.6 – 28%) were less common or no different. Adolescents in 2015 were spending less time sleeping, had higher BMIs and a greater proportion perceived themselves as overweight.

**Conclusion:** Given health-related behaviours are often cited as risk-factors for poor mental health, our findings suggest relationships between these factors might be more complex and dynamic in nature than currently understood. Striking increases in mental health difficulties, BMI and poor sleep related behaviours highlight an increasing public health challenge.

## Introduction

The focus on adolescent health has been increasing in recent years,^1^ with a growing recognition that these years are pivotal in the development and maintenance of health behaviours and outcomes through the lifecourse.^2,3^ Adolescence is a key period for mental health disorder onset with half of lifetime onset by age 14.^4^ Research over previous decades suggests that the prevalences of mental health problems are increasing in UK teenagers,^5,6^ which is mirrored in studies across different countries.^7,8^ An international systematic review investigating secular trends in adolescent mental health from the previous century into the start of this century concluded that internalizing symptoms seem to be increasing, finding more consistent evidence for increases in girls compared to boys.^9^ Most studies in this review focussed on internalizing symptoms or general psychological distress, making conclusions about externalising behaviours less possible. There are few studies comparing changing trends in the millennial generations, and prevalence studies suggest that mental health problems in mid-adolescence might have increased even further in recent years.^10^

In contrast, while prevalence of internalizing mental health problems seems to be increasing, young people in the UK are becoming less likely to be underage substance users. Office for National Statistics reports collected from secondary school pupils in England have found prevalence of alcohol use, smoking, cannabis use and other drug use among 14 year old pupils have consistently fallen since 1982 when the survey was first undertaken.^11^ For example, in 1982 16% of 14 year olds described themselves as regular smokers. In 2014 this had fallen to just 4%, and the drop was consistent across genders. This decrease in use has become particularly pronounced since the early 2000s.^11^

Given that various health behaviours including but not limited to substance use are implicated in risk for poor mental health,^12-17^ investigating the relationships between these secular trends is important to explore causal relationships, the aetiology of mental ill-health, and potentially to inform interventions to try and reverse the increasing prevalence of mental health problems. It is therefore surprising that to date there has been little attempt to combine investigations of secular trends in mental health with changes in health related behaviours. We also know little about trends in other health related behaviours such as sleep, risky sexual behaviour, body satisfaction and physical activity that might also be causal risk factors for mental ill-health and substance use.^12-17^

In the current study we use two cohorts of UK adolescents born a decade apart (1991/92 and 2000/02) in order to identify secular trends in mental health, considering both internalizing and externalizing symptoms, and a number of related health behaviours including substance use, sleep behaviours, weight, physical and sexual activity. In particular we attempt to make the variables and datasets as comparable as possible by harmonizing the variables and performing two different techniques (propensity score matching and entropy balancing) to increase the comparability of the cohorts. The prevalence of a number of these behaviours differ between males and females, and some studies report different trends in males and females.^6^ We therefore also empirically examine sex differences in time trends in these outcomes.

## Method

### Participants

***Avon Longitudinal Study of Parents and Children (ALSPAC)*** is a cohort born in 1991-92. ALSPAC recruited 14,541 pregnant women resident in Avon, UK with expected dates of delivery 1st April 1991 to 31st December 1992. When the oldest children were approximately 7 years of age, an attempt was made to bolster the initial sample with eligible cases who had failed to join the study originally. The total sample size for analyses using any data collected after the age of seven is therefore 15,247 pregnancies, resulting in 15,458 foetuses. Of this total sample of 15,458 foetuses, 14,775 were live births and 14,701 were alive at 1 year of age.^18,19^ The study website contains details of all the data that is available through a fully searchable data dictionary and variable search tool (http://www.bristol.ac.uk/alspac/researchers/our-data/). Ethics approval for the study was obtained from the ALSPAC Ethics and Law Committee and the Local Research Ethics Committees. Data were collected frequently via different modalities, with clinic visits and postal questionnaires having taken place in adolescence every year. This study uses data from ages 13, 14 and 15. In the current study, data were available from 6132 participants at age 14 representing 41.7% of the 14701 participants alive past 1 year. Attrition is predicted by a range of variables in ALSPAC including lower educational level, male gender, non-White ethnicity, and eligibility for free school meals.^18^

***Millennium Cohort Study (MCS)*** is a cohort of 19,517 children born in 2000-02 sampled from the whole of the UK.^20^ Data so far have been collected in 6 sweeps at ages 9 months, 3, 5, 7, 11 and 14 years. The <u>study website</u> (http://www.cls.ioe.ac.uk/) contains details regarding all the data available and information on accessing the datasets. Ethics approval for the age 14 sweep was obtained from the National Research Ethics Service Research Ethics Committee. At the age 14 sweep, 15415 families were issued into the field (those not issued due to emigration, permanent refusal, untraceability), of which 11726 families participated in the age 14 sweep (representing 60.9% of the original sample).^21^Attrition at the age 14 sweep compared to the full sample is predicted by a range of demographic variables including male gender, Black ethnicity, lower occupational and educational level and single parent family.^22^

For this study, we analysed data from participants who had provided data on at least one of the outcome variables at the age 14 sweeps of the studies (depressive symptoms, smoking, alcohol, cannabis and other drugs; ALSPAC N= 6132, MCS N= 11351). Furthermore, participants without the demographic data required for increasing the comparability of the datasets (sex, ethnicity, age, maternal education and maternal age) were excluded, resulting in an analysis sample of 5627 from ALSPAC and 11318 from MCS.

There have been changes in socio-demographic characteristics of the country in the ten years between these cohorts (e.g. higher proportion ethnic minorities, higher education levels) and in addition the two cohorts represent different regions that might have different characteristics with ALPSAC being a regional and MCS a national cohort. For instance, around one-fifth of the MCS sample are ethnic minorities compared to around 4% in ALSPAC. To minimise the bias in time trends socio-demographic differences in the sample might cause, we control for socio-demographic factors in analysis and in addition employ two additional approaches (propensity score matching and entropy balancing) to increase the comparability of the cohorts.

### Measures

The measures used in this study (Table 1) include socio-demographic indicators (used for increasing cohort comparability), mental ill-health (depressive symptoms, self-harm, parent reported difficulties), substance use (alcohol, smoking, cannabis and other drugs), antisocial behaviours (assault, graffiti, vandalism, shoplifting and rowdy behaviour) and other health related behaviours (including sleep, weight, weight perception and sexual activity). In a few instances (self-harm, sleep behaviours, parent-rated difficulties), the variables of interest were not available in ALSPAC at age 14, but available in the sweep immediately before (age 13) or afterwards (age 15) and where this is the case is clearly indicated in the table. Table 1 also presents the details of the harmonised variables that were subsequently used in analysis. Some of the variables were more readily comparable than others, for instance both studies used the Short Moods and Feelings Questionnaire^23^ to assess depressive symptoms, parent-rated Strengths and Difficulties Questionnaire^24^ to measure difficulties and the same set of questions to record sexual activity. Other variables were harmonised through a process of creating new, comparable variables across the datasets (e.g. alcohol, smoking). For self-harm, even after harmonisation the resulting variable is not truly comparable due to different time scales of the question asked, which needs to be borne in mind when interpreting findings. Lastly, some health related behaviours that we planned to harmonise and investigate (physical activity) were substantially differently measured and harmonised measures could not be derived.

**Table 1.** Measures in ALSPAC and MCS for each domain and the harmonised variable

| Outcome | ALSPAC (2005) | MCS (2015) | Harmonised variable |
| --- | --- | --- | --- |
| Depressive symptoms | <sup>a</sup> 13-item Short Moods and Feelings Questionnaire | <sup>b</sup> 13-item Short Moods and Feelings Questionnaire | Total depressive symptoms score (continuous)<br>yes/no variable based on clinical threshold $\geq 12$ |
| Self-harm | <sup>a</sup> (Measured at age 15)<br>Over your whole lifetime have you ever tried to harm yourself or kill yourself? | <sup>b</sup> In the past year have you hurt yourself on purpose in any way? | 0 Not self harmed<br>1 Have self harmed<br><u>NOTE:</u> ALSPAC is lifetime but MCS is in past year |
| Antisocial behaviours | <sup>b</sup> How often in the last year have you done any of the following? (categorical)<br>- Not at all<br>- Just once<br>- 3-5 times<br>- 6+ times<br>Hit kicked or punched someone on purpose<br>Been rowdy or rude in a public place so that people complained or you got in to trouble<br>Written things or sprayed paint on a property that did not belong to you<br>Deliberately damaged or destroyed property that did not belong to you<br>Taken something from a shop without paying for it<br>Other items included skipping school, breaking and stealing from various different places, carrying a knife or weapon for protection, setting fire, stealing a vehicle, not paying correct fare on public transport, using force to steal. | <sup>b</sup> In the last 12 months have you: (yes/no)<br>Pushed or shoved /hit/slapped/punched someone?<br>Been noisy or rude in a public place so that people complained or got you into trouble?<br>Written things or spray painted on a building, fence or train or anywhere else where you shouldn't have?<br>On purpose damaged anything in a public place that didn't belong to you, for example by burning, smashing or breaking things like cars, bus shelters and rubbish bins?<br>Taken something from a shop without paying for it?<br>Other items included using a weapon on someone, stealing from someone, hacking into computers, sending computer viruses. | 0 No<br>1 Yes<br>Assault<br>Rowdy behaviour<br>Graffiti<br>Vandalism<br>Shoplifting |
| Parent rated difficulties | <sup>c</sup> (Measured at age 13)<br>Strengths and difficulties Questionnaire | <sup>c</sup> Strengths and difficulties Questionnaire | 5 continuous subscale scores<br>5 yes/no variables based on the 'abnormal' cutoff<br>Subscales: emotional symptom, conduct problems, hyperactivity, peer problems, prosocial behaviour |
| Alcohol use | <sup>a</sup> Have you ever tried alcohol with/without your parents' permission? (yes/no)<br>What is the most alcoholic drinks you've had in a single evening? (continuous variable)<br>How many times have you done this in the last year? (continuous variable) | <sup>b</sup> Have you ever had an alcoholic drink? That is more than a few sips. (yes/no)<br>How many times have you had an alcoholic drink in the last 12 months? (7 response options from never to over 40) | 0 Never drank a whole drink<br>1 Nothing in last 12 months/never drank 5 or more<br>2 1-2 times drank 5 or more alcoholic drinks in one evening<br>3 3 or more times drank 5 or more alcoholic drinks in one evening |
|  |  | Have you ever had five or more alcoholic drinks at a time? A drink is half a pint of lager, beer or cider, one alcopop, a small glass of wine, or a measure of spirits. (yes/no) |  |
|  |  | How many times have you had five or more alcoholic drinks at a time in the last 12 months?<br>- Never<br>- 1-2 times<br>- 3-5 times<br>- 6-9 times<br>- 10 or more time |  |
| Smoking frequency | <sup>a</sup> Frequency teenager has smoked cigarettes in the past 6 months:<br>- 1-3 times;<br>- >4 times;<br>- once per week;<br>- never.<br>Number of cigarettes smoked per week in the last 6 months for weekly users (continuous variable) | <sup>b</sup> Please read the following statements carefully and decide which one best describes you. Do not include electronic cigarettes (e-cigarettes).<br>- I have never smoked cigarettes;<br>- I have only ever tried smoking cigarettes once;<br>- I used to smoke sometimes but I never smoke a cigarette now;<br>- I sometimes smoke cigarettes now but I don't smoke as many as one a week;<br>- I usually smoke between one and six cigarettes a week;<br>- I usually smoke more than six cigarettes a week. | 0 Non smoker<br>1 Occasional smoker, not weekly<br>2 Smokes 1-6 cigarettes a week<br>3 Smokes more than 6 cigarettes a week |
| Cannabis | <sup>a</sup> Have you ever used cannabis? (yes/no) | <sup>b</sup> Have you ever tried any of the following things? Cannabis (also known as weed, marijuana, dope, hash or skunk)? (yes/no) | 0 Never used cannabis<br>1 Have tried cannabis |
| Other drugs | <sup>a</sup> Teenager has been offered drugs? (yes/no)<br><if yes to above> Teenager has used drugs other than cannabis to feel good/get high? (yes/no) | <sup>b</sup> Have you ever tried any of the following things? Any other illegal drug (such as ecstasy, cocaine, speed)? (yes/no) | 0 Never used other drugs<br>1 Have tried other drugs |
| BMI | <sup>a</sup> Height and weight measured by interviewer | <sup>a</sup> Height and weight measured by interviewer | BMI derived from height and weight<br>Obese (0=no, 1=yes) derived using IOTF threshold |
| Weight perception | <sup>b</sup> How do you describe your weight?<br>- Very underweight<br>- Slightly underweight<br>- About the right weight<br>- Slightly overweight<br>- Very overweight | <sup>b</sup> Which of these do you think you are?<br>- Underweight<br>- About the right weight<br>- Slightly overweight<br>- Very overweight | Perceive themselves:<br>1 Underweight<br>2 About the right weight<br>3 Slightly overweight<br>4 Very overweight |
| Sleep | <sup>b</sup> (Measured at age 15)<br>What time do you usually get in to be on school days? (continuous - reported in hours and minutes am/pm)<br>What time do you usually get into bed on weekend days? | <sup>b</sup> About what time do you usually go to sleep on a school night?<br>About what time do you usually go to sleep on the nights when you do not have school the next day?<br>- Before 9pm | 4 categorical sleep variables:<br>Schoolday bedtime<br>Non schoolday bedtime<br>1 Before 9pm<br>2 9 - 9.59pm<br>3 10 - 10.59pm |
|  | (continuous - reported in hours and minutes am/pm) | - 9 - 9.59pm<br>- 10 - 10.59pm<br>- 11pm - midnight<br>- After midnight | 4 11pm - midnight<br>5 After midnight<br>(11 pm and later classified as late bedtime) |
|  | What time do you usually wake up on school days?<br>(continuous - reported in hours and minutes am/pm) | About what time do you usually wake up on a school day?<br>- Before 6am | Schoolday waketime<br>1 Before 6am |
|  | What time do you usually wake up on weekend days?<br>(continuous - reported in hours and minutes am/pm) | - 6 - 6.59am<br>- 7 - 7.59am<br>- 8 - 8.59am<br>- 9am or later | 2 6 - 6.59am<br>3 7 - 7.59am<br>4 8 - 8.59am<br>5 9am or later<br>(before 7 am classified as early waketime) |
|  |  | About what time do you wake up in the morning on the days when you do not have school?<br>- Before 8am<br>- 8 - 8.59am<br>- 9 - 9.59am<br>- 10 - 10.59am<br>- 11 - 11.59am<br>- Midday or later | Non schoolday waketime<br>1 Before 8am<br>2 8 - 8.59am<br>3 9 - 9.59am<br>4 10 - 10.59am<br>5 11 - 11.59am<br>6 Midday or later<br>(before 9 am classified as early waketime) |
| Sexual intercourse | <sup>b</sup> Series of questions regarding intimate contact with someone else leading to:<br>Have you had sexual intercourse with another person in the past year? | <sup>b</sup> Series of questions regarding intimate contact with someone else leading to:<br>In the last 12 months have you had sexual intercourse with another young person? | 0 have not had sex<br>1 had sex |
| Physical activity | <sup>b</sup> Frequency data available on approx. 80 specific activities (riding a bike, skipping, gardening, walking the dog, cricket etc) | <sup>b</sup> On how many days in the last week did you do a total of at least an hour of moderate to vigorous physical activity? By moderate to vigorous we mean any physical activity that makes you get warmer, breathe harder and makes your heart beat faster, e.g. riding a bike, running, playing football, swimming, dancing, etc.<br>- Every day<br>- 5-6 days<br>- 3-4 days<br>- 1-2 days<br>- Not at all | Not harmonised |

<sup>a</sup> Measured via interview; <sup>b</sup> Measured via self completion; <sup>c</sup>Parent reported

## Analysis

### Increasing comparability of the datasets

To increase comparability of the samples by accounting for key socio-demographic differences between these samples, participants from the larger MCS sample were matched or weighted to make them comparable to the ALSPAC sample on key demographic factors including sex, age, ethnicity, maternal education and maternal age at birth. This was done using two approaches: propensity score matching^25^ and entropy balancing.^26^ Both approaches aim to reduce the probability that differences between samples on outcomes of interest are because of sample differences on relevant demographic variables.^26^ Table S1 shows the differences in these characteristics in the samples before and after these procedures were applied.

*Propensity score matching* is based on a propensity score, which is derived from weighting schemes based on the criteria that are to be matched to identify individuals from the larger, control group that are most like each of the individuals in the treatment group (in this case ALSPAC) across a range of variables as specified. Propensity score matching was conducted in STATA using psmatch2.^27^

*Entropy balancing* is a multivariate reweighting method that calibrates unit weights such that two samples are balanced on a range of pre-specified variables, hence increasing comparability for the estimation of treatment, or in this case cohort, effects.^26^ The application of this approach creates an entropy balancing weight value for all participants in the MCS sample, which is then used as a weight when estimating prevalences in the MCS sample. This approach allows the utilisation of the full available MCS cohort, instead of selecting a matched sub-sample like the propensity score matching approach. Entropy balancing was conducted in STATA using ebalance.^28^

### Missing data

In ALSPAC 15.6% of the total cells were missing (ranging from <1% for substance use, 1.8% for mental health, ∼24% for antisocial behaviours and ∼26% for sleep behaviours). In the MCS samples, 1% of cells were missing in the MCS propensity matched sample and 1.2% in the full MCS sample. Multiple imputations (20 imputations) were carried out using chained equations separately in the two cohorts.

### Estimating cohort differences

Four estimates (ALSPAC, MCS nationally representative, MCS propensity matched and MCS entropy balanced) of the prevalences and descriptive statistics (means and % with 95% CIs) for each of the harmonised outcome variables were first estimated. In addition, we estimated odds ratios (Figure 1) of the cohort effects (MCS compared to ALSPAC) for the prevalence of high mental health or risky health behaviours using logistic regressions. Lastly, we also examined sex into cohort effect interactions with the *ebalancing* weight to examine whether any outcomes have trends that are different rates in males and females.

**Figure 1.**
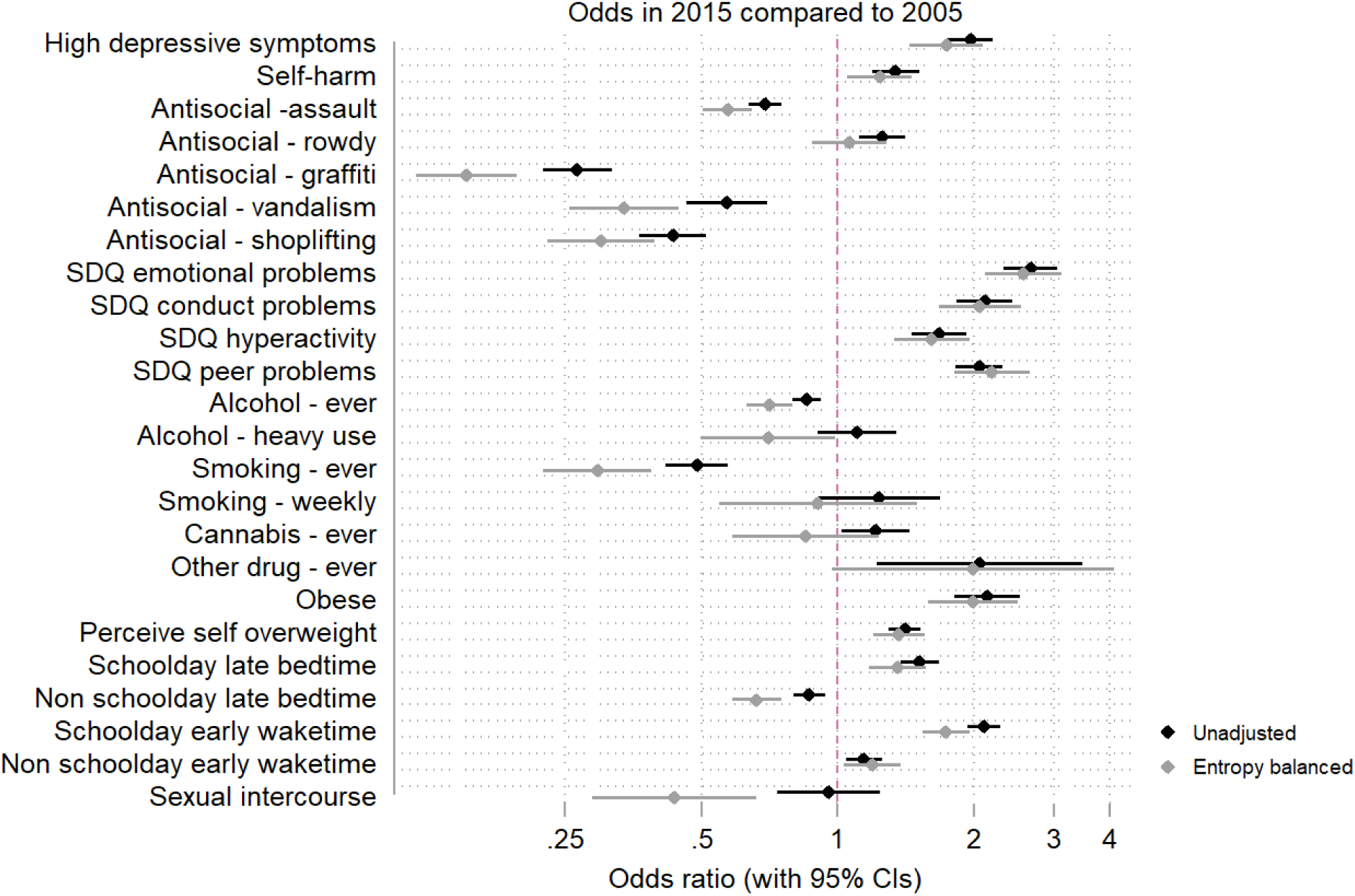
ORs (95% CI) for poorer outcomes in 2015 (MCS) vs. 2005 (ALSPAC). Unadjusted estimates and estimates using entropy balancing weights are both presented.

For ease of interpretation for the reader, throughout the rest of the paper we refer to the ALSPAC variables year of collection as 2005 and the MCS variables as 2015.

## Results

There were no differences between the samples in sex distribution and maternal age at birth (Table S1). Regarding the other characteristics, as expected, MCS had higher proportions of ethnic minorities and higher levels of maternal higher education. Of the two approaches used to increase comparability, the propensity matching resulted in the two samples becoming more similar, for example ethnic minorities were less than 4% in the ALSPAC cohort compared to more than 20% in the full MCS cohort, while the propensity matched MCS cohort consisted of around 10% of minority ethnic individuals. In contrast, the entropy balancing (based on the generated entropy weights) resulted in matched estimates across demographic characteristics in the two cohorts.

Estimates from the propensity score matched sample and the entropy balancing in the MCS were very similar in most cases and for most outcomes, different from the MCS nationally representative estimates, indicating the relevance of adjusting the estimates when estimating cohort differences. The descriptive statistics indicated that there were more young people with mental health problems, as indicated by greater proportion above depression threshold and reporting self-harming, in 2015 compared to 2005 (but note the self-harm behaviour question was limited to past 12 months in 2015 compared to lifetime in 2005). Antisocial behaviour and substance use rates were lower in 2015 compared to 2005. Parent reported difficulties highlighted higher rates of emotional, conduct and hyperactivity symptoms and greater levels of problems getting along with peers in 2015 compared to 2005. With regards to the health-related behaviours, the more recent cohort had a higher BMI on average and larger numbers also perceived themselves to being overweight. The data on sleep behaviours indicated that on weekdays young people in 2015 were more likely to sleep later and more likely to wake up earlier. Weekend sleep and wake times were more similar between the cohorts. A greater proportion of adolescents in 2005 reported having had sexual intercourse by this age compared to in 2015. Due to the higher comparability and complete sample size using entropy weights and the similar estimates produced with entropy and propensity adjustment, entropy balancing is used for subsequent regression analyses comparing the two cohorts and the sex by cohort interactions.

Figure 1 illustrates odds of outcomes in the MCS sample (2015) compared to the ALSPAC sample (2005) using both a direct comparison approach and using estimates applying the entropy balancing weights. Estimates were similar for most of the mental health and some health related behaviour outcomes based on the two approaches, but there was some noticeable upward or downward bias for some outcomes, for instance with entropy balancing the lower odds in 2015 compared to 2005 are more stark for antisocial and risky health behaviours; highlighting the potential relevance of using methods to increase the comparability of cohorts when estimating cohort differences and trends.

Descriptives stratified by sex are presented in Table 4. Depressive symptoms, self-harm, overweight perception were higher in females and antisocial behaviours, peer problems higher in males. Regression analysis with the entropy balancing weight were estimated to examine sex-by-cohort interactions. Most health-related behaviours showed little or no sex differences in prevalence. There were no sex-by-cohort interactions for most of the variables included in this study, indicating that rates of change or increased/decreased odds were similar in males and females. There was evidence of sex differences in cohort effects for some antisocial behaviours (e.g. assault OR_male_ = 0.66, OR _female_ = 0.45), parent-reported conduct problems (OR_male_ = 2.74, OR _female_ = 1.38) and having tried alcohol (OR_male_ = 0.85, OR _female_= 0.59), where odds of these behaviours in 2015 compared to 2005 were lower in females compared to males (irrespective of whether overall odds were lower or higher in 2015). Odds ratios separately by sex were estimated and presented in Figure 2.

**Table 2.**
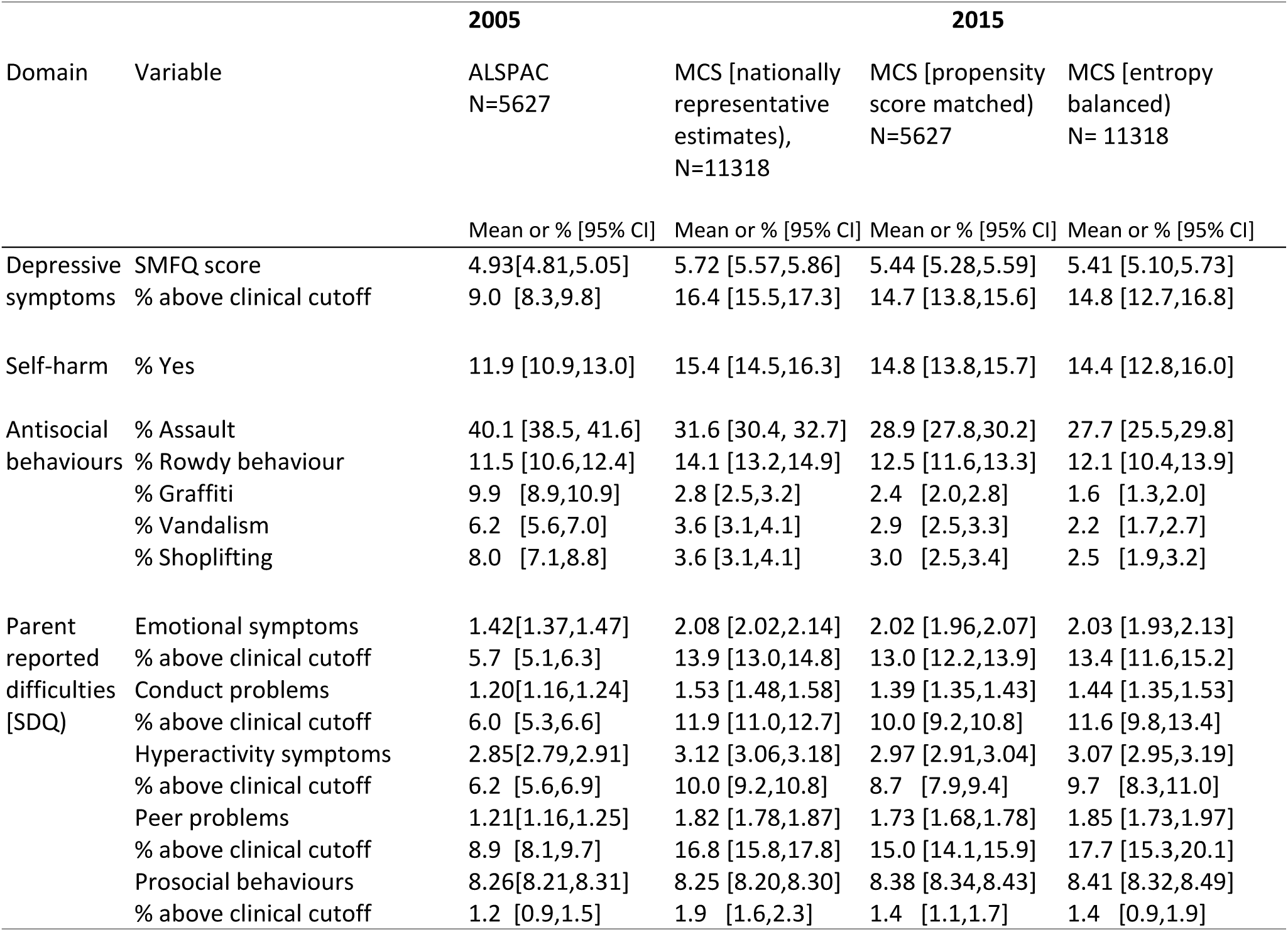
Descriptive statistics for mental health outcomes in 2005 (ALSPAC) and 2015 (MCS)

**Table 3.**
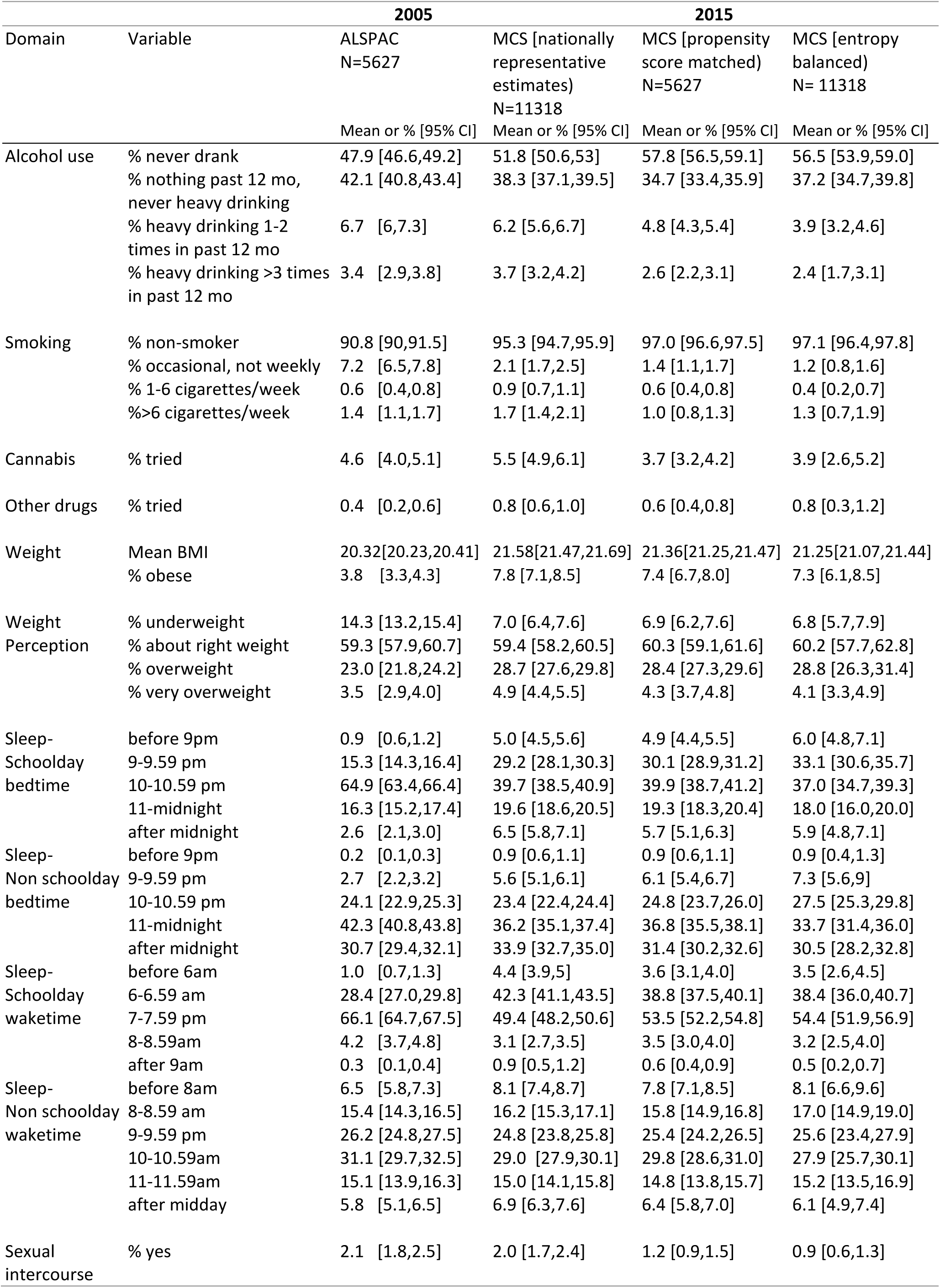
Descriptive statistics for health-related behaviours in 2005 (ALSPAC) and 2015 (MCS)

**Table 4.**
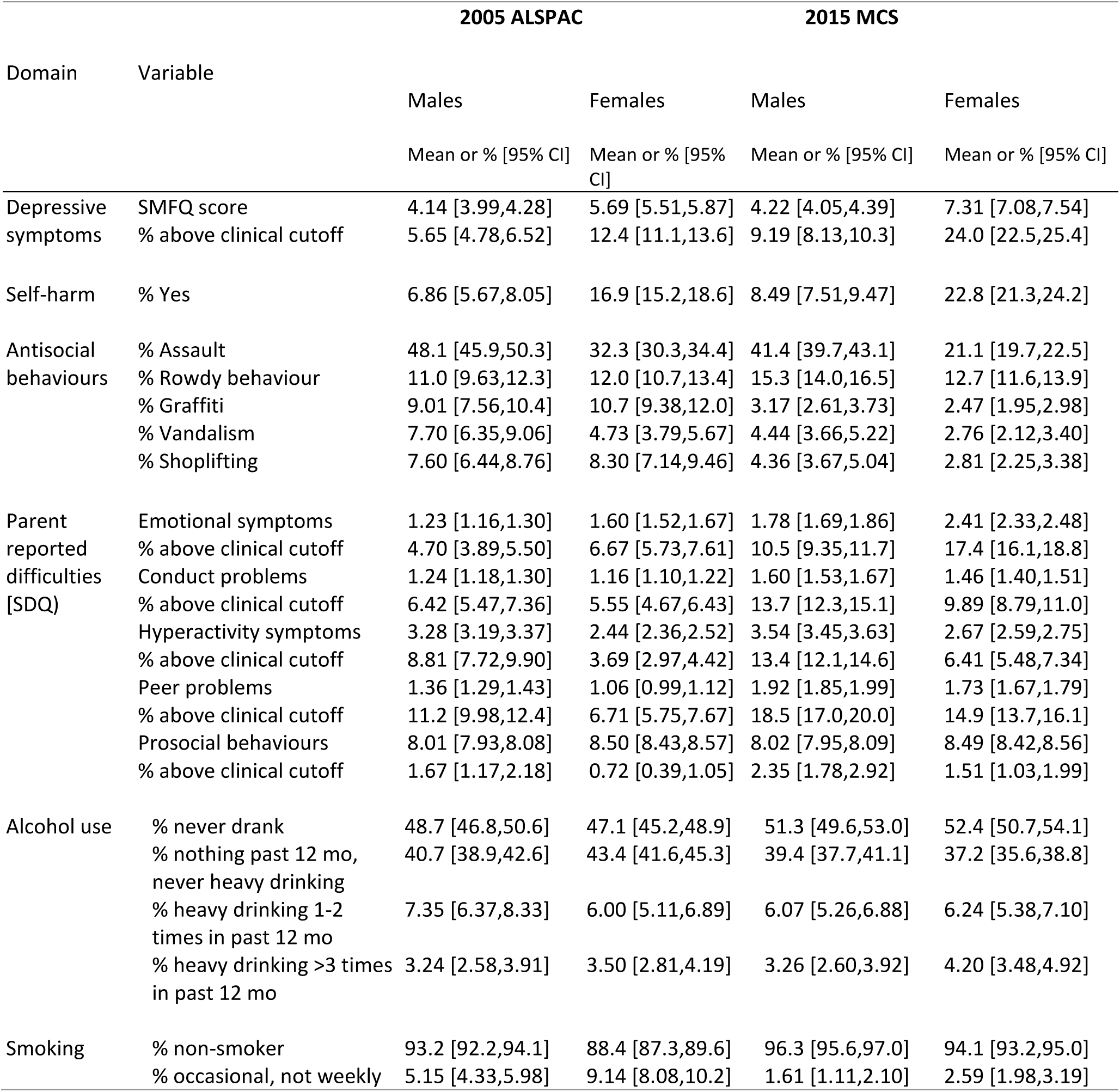
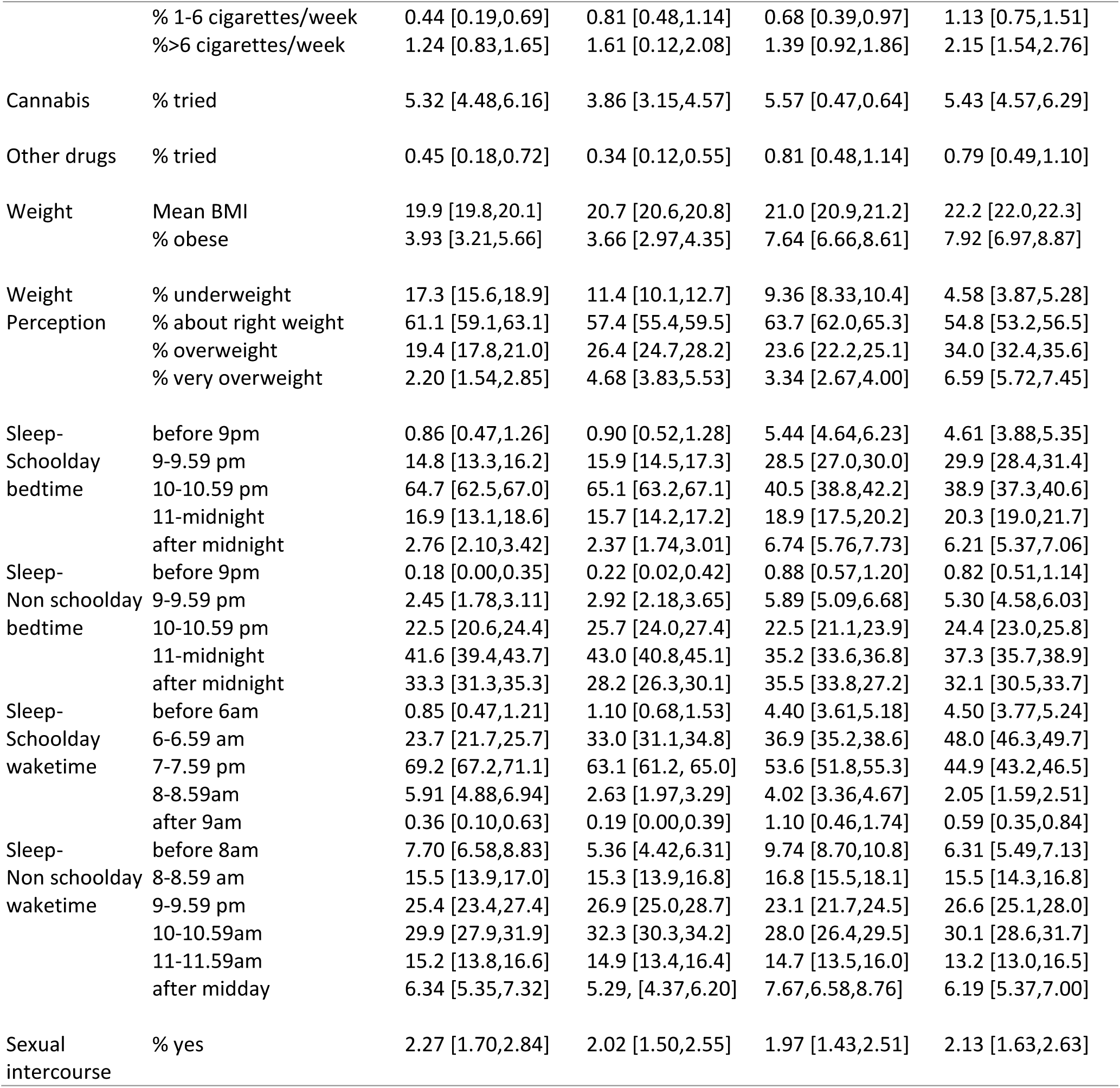
Descriptive statistics for mental health and health related behaviours in 2005 (ALSPAC) and 2015 (MCS) by sex

| Domain | Variable | 2005 ALSPAC |  | 2015 MCS |  |
| --- | --- | --- | --- | --- | --- |
|  |  | Males | Females | Males | Females |
|  |  | Mean or % [95% CI] | Mean or % [95% CI] | Mean or % [95% CI] | Mean or % [95% CI] |
| Depressive symptoms | SMFQ score | 4.14 [3.99,4.28] | 5.69 [5.51,5.87] | 4.22 [4.05,4.39] | 7.31 [7.08,7.54] |
|  | % above clinical cutoff | 5.65 [4.78,6.52] | 12.4 [11.1,13.6] | 9.19 [8.13,10.3] | 24.0 [22.5,25.4] |
| Self-harm | % Yes | 6.86 [5.67,8.05] | 16.9 [15.2,18.6] | 8.49 [7.51,9.47] | 22.8 [21.3,24.2] |
| Antisocial behaviours | % Assault | 48.1 [45.9,50.3] | 32.3 [30.3,34.4] | 41.4 [39.7,43.1] | 21.1 [19.7,22.5] |
|  | % Rowdy behaviour | 11.0 [9.63,12.3] | 12.0 [10.7,13.4] | 15.3 [14.0,16.5] | 12.7 [11.6,13.9] |
|  | % Graffiti | 9.01 [7.56,10.4] | 10.7 [9.38,12.0] | 3.17 [2.61,3.73] | 2.47 [1.95,2.98] |
|  | % Vandalism | 7.70 [6.35,9.06] | 4.73 [3.79,5.67] | 4.44 [3.66,5.22] | 2.76 [2.12,3.40] |
|  | % Shoplifting | 7.60 [6.44,8.76] | 8.30 [7.14,9.46] | 4.36 [3.67,5.04] | 2.81 [2.25,3.38] |
| Parent reported difficulties [SDQ] | Emotional symptoms | 1.23 [1.16,1.30] | 1.60 [1.52,1.67] | 1.78 [1.69,1.86] | 2.41 [2.33,2.48] |
|  | % above clinical cutoff | 4.70 [3.89,5.50] | 6.67 [5.73,7.61] | 10.5 [9.35,11.7] | 17.4 [16.1,18.8] |
|  | Conduct problems | 1.24 [1.18,1.30] | 1.16 [1.10,1.22] | 1.60 [1.53,1.67] | 1.46 [1.40,1.51] |
|  | % above clinical cutoff | 6.42 [5.47,7.36] | 5.55 [4.67,6.43] | 13.7 [12.3,15.1] | 9.89 [8.79,11.0] |
|  | Hyperactivity symptoms | 3.28 [3.19,3.37] | 2.44 [2.36,2.52] | 3.54 [3.45,3.63] | 2.67 [2.59,2.75] |
|  | % above clinical cutoff | 8.81 [7.72,9.90] | 3.69 [2.97,4.42] | 13.4 [12.1,14.6] | 6.41 [5.48,7.34] |
|  | Peer problems | 1.36 [1.29,1.43] | 1.06 [0.99,1.12] | 1.92 [1.85,1.99] | 1.73 [1.67,1.79] |
|  | % above clinical cutoff | 11.2 [9.98,12.4] | 6.71 [5.75,7.67] | 18.5 [17.0,20.0] | 14.9 [13.7,16.1] |
|  | Prosocial behaviours | 8.01 [7.93,8.08] | 8.50 [8.43,8.57] | 8.02 [7.95,8.09] | 8.49 [8.42,8.56] |
|  | % above clinical cutoff | 1.67 [1.17,2.18] | 0.72 [0.39,1.05] | 2.35 [1.78,2.92] | 1.51 [1.03,1.99] |
| Alcohol use | % never drank | 48.7 [46.8,50.6] | 47.1 [45.2,48.9] | 51.3 [49.6,53.0] | 52.4 [50.7,54.1] |
|  | % nothing past 12 mo, never heavy drinking | 40.7 [38.9,42.6] | 43.4 [41.6,45.3] | 39.4 [37.7,41.1] | 37.2 [35.6,38.8] |
|  | % heavy drinking 1-2 times in past 12 mo | 7.35 [6.37,8.33] | 6.00 [5.11,6.89] | 6.07 [5.26,6.88] | 6.24 [5.38,7.10] |
|  | % heavy drinking >3 times in past 12 mo | 3.24 [2.58,3.91] | 3.50 [2.81,4.19] | 3.26 [2.60,3.92] | 4.20 [3.48,4.92] |
| Smoking | % non-smoker | 93.2 [92.2,94.1] | 88.4 [87.3,89.6] | 96.3 [95.6,97.0] | 94.1 [93.2,95.0] |
|  | % occasional, not weekly | 5.15 [4.33,5.98] | 9.14 [8.08,10.2] | 1.61 [1.11,2.10] | 2.59 [1.98,3.19] |
|  | % 1-6 cigarettes/week | 0.44 [0.19,0.69] | 0.81 [0.48,1.14] | 0.68 [0.39,0.97] | 1.13 [0.75,1.51] |
|  | %>6 cigarettes/week | 1.24 [0.83,1.65] | 1.61 [0.12,2.08] | 1.39 [0.92,1.86] | 2.15 [1.54,2.76] |
| Cannabis | % tried | 5.32 [4.48,6.16] | 3.86 [3.15,4.57] | 5.57 [0.47,0.64] | 5.43 [4.57,6.29] |
| Other drugs | % tried | 0.45 [0.18,0.72] | 0.34 [0.12,0.55] | 0.81 [0.48,1.14] | 0.79 [0.49,1.10] |
| Weight | Mean BMI | 19.9 [19.8,20.1] | 20.7 [20.6,20.8] | 21.0 [20.9,21.2] | 22.2 [22.0,22.3] |
|  | % obese | 3.93 [3.21,5.66] | 3.66 [2.97,4.35] | 7.64 [6.66,8.61] | 7.92 [6.97,8.87] |
| Weight Perception | % underweight | 17.3 [15.6,18.9] | 11.4 [10.1,12.7] | 9.36 [8.33,10.4] | 4.58 [3.87,5.28] |
|  | % about right weight | 61.1 [59.1,63.1] | 57.4 [55.4,59.5] | 63.7 [62.0,65.3] | 54.8 [53.2,56.5] |
|  | % overweight | 19.4 [17.8,21.0] | 26.4 [24.7,28.2] | 23.6 [22.2,25.1] | 34.0 [32.4,35.6] |
|  | % very overweight | 2.20 [1.54,2.85] | 4.68 [3.83,5.53] | 3.34 [2.67,4.00] | 6.59 [5.72,7.45] |
| Sleep-Schoolday bedtime | before 9pm | 0.86 [0.47,1.26] | 0.90 [0.52,1.28] | 5.44 [4.64,6.23] | 4.61 [3.88,5.35] |
|  | 9-9.59 pm | 14.8 [13.3,16.2] | 15.9 [14.5,17.3] | 28.5 [27.0,30.0] | 29.9 [28.4,31.4] |
|  | 10-10.59 pm | 64.7 [62.5,67.0] | 65.1 [63.2,67.1] | 40.5 [38.8,42.2] | 38.9 [37.3,40.6] |
|  | 11-midnight | 16.9 [13.1,18.6] | 15.7 [14.2,17.2] | 18.9 [17.5,20.2] | 20.3 [19.0,21.7] |
|  | after midnight | 2.76 [2.10,3.42] | 2.37 [1.74,3.01] | 6.74 [5.76,7.73] | 6.21 [5.37,7.06] |
| Sleep-Non schoolday bedtime | before 9pm | 0.18 [0.00,0.35] | 0.22 [0.02,0.42] | 0.88 [0.57,1.20] | 0.82 [0.51,1.14] |
|  | 9-9.59 pm | 2.45 [1.78,3.11] | 2.92 [2.18,3.65] | 5.89 [5.09,6.68] | 5.30 [4.58,6.03] |
|  | 10-10.59 pm | 22.5 [20.6,24.4] | 25.7 [24.0,27.4] | 22.5 [21.1,23.9] | 24.4 [23.0,25.8] |
|  | 11-midnight | 41.6 [39.4,43.7] | 43.0 [40.8,45.1] | 35.2 [33.6,36.8] | 37.3 [35.7,38.9] |
|  | after midnight | 33.3 [31.3,35.3] | 28.2 [26.3,30.1] | 35.5 [33.8,27.2] | 32.1 [30.5,33.7] |
| Sleep-Schoolday waketime | before 6am | 0.85 [0.47,1.21] | 1.10 [0.68,1.53] | 4.40 [3.61,5.18] | 4.50 [3.77,5.24] |
|  | 6-6.59 am | 23.7 [21.7,25.7] | 33.0 [31.1,34.8] | 36.9 [35.2,38.6] | 48.0 [46.3,49.7] |
|  | 7-7.59 pm | 69.2 [67.2,71.1] | 63.1 [61.2, 65.0] | 53.6 [51.8,55.3] | 44.9 [43.2,46.5] |
|  | 8-8.59am | 5.91 [4.88,6.94] | 2.63 [1.97,3.29] | 4.02 [3.36,4.67] | 2.05 [1.59,2.51] |
|  | after 9am | 0.36 [0.10,0.63] | 0.19 [0.00,0.39] | 1.10 [0.46,1.74] | 0.59 [0.35,0.84] |
| Sleep-Non schoolday waketime | before 8am | 7.70 [6.58,8.83] | 5.36 [4.42,6.31] | 9.74 [8.70,10.8] | 6.31 [5.49,7.13] |
|  | 8-8.59 am | 15.5 [13.9,17.0] | 15.3 [13.9,16.8] | 16.8 [15.5,18.1] | 15.5 [14.3,16.8] |
|  | 9-9.59 pm | 25.4 [23.4,27.4] | 26.9 [25.0,28.7] | 23.1 [21.7,24.5] | 26.6 [25.1,28.0] |
|  | 10-10.59am | 29.9 [27.9,31.9] | 32.3 [30.3,34.2] | 28.0 [26.4,29.5] | 30.1 [28.6,31.7] |
|  | 11-11.59am | 15.2 [13.8,16.6] | 14.9 [13.4,16.4] | 14.7 [13.5,16.0] | 13.2 [13.0,16.5] |
|  | after midday | 6.34 [5.35,7.32] | 5.29, [4.37,6.20] | 7.67,6.58,8.76] | 6.19 [5.37,7.00] |
| Sexual intercourse | % yes | 2.27 [1.70,2.84] | 2.02 [1.50,2.55] | 1.97 [1.43,2.51] | 2.13 [1.63,2.63] |

**Figure 2.**
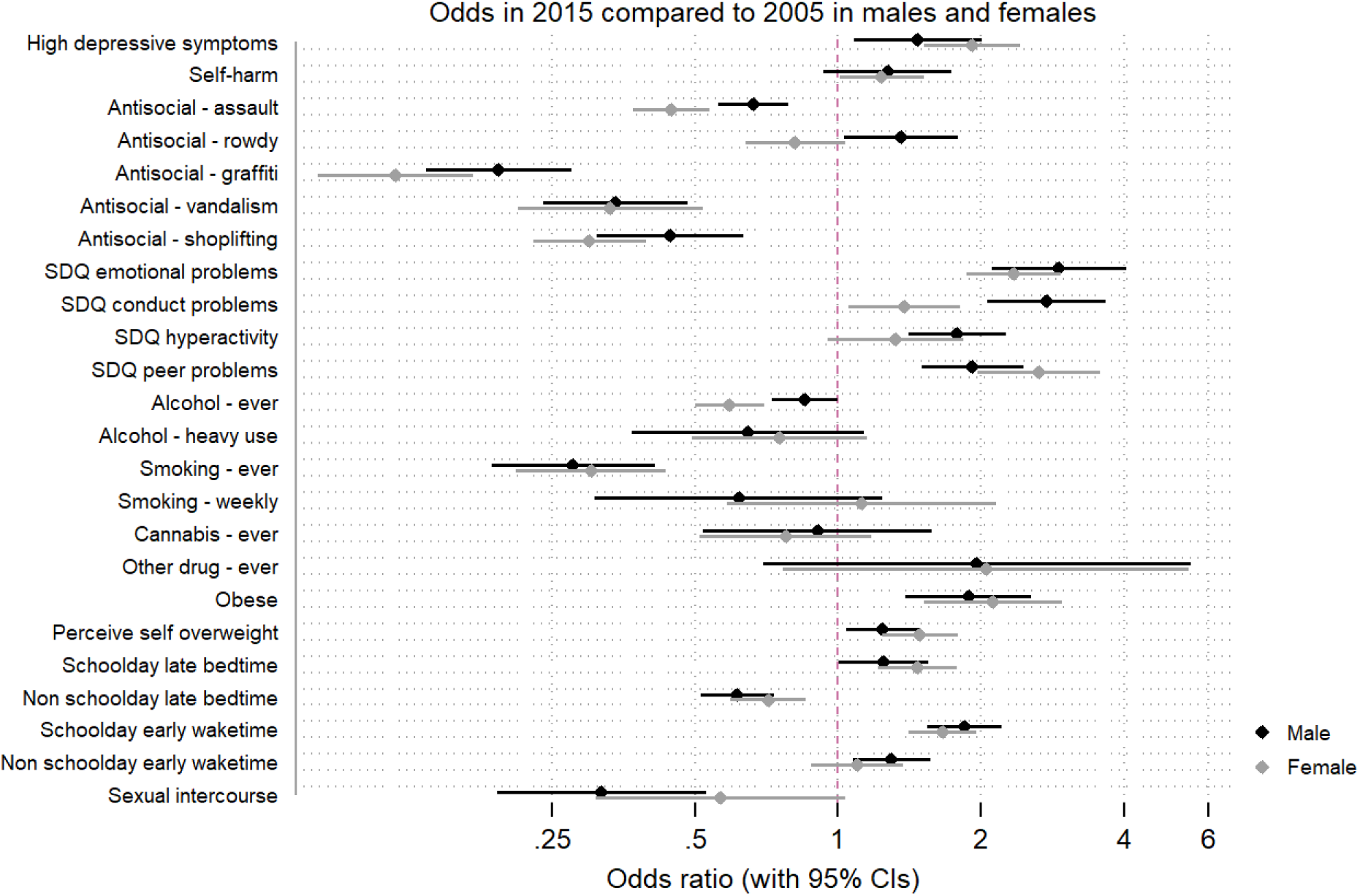
ORs (95% CI) for poorer outcomes in 2015 (MCS) vs. 2005 (ALSPAC) for males and females.

## Discussion

The current study examined recent trends in a range of mental health and health related behaviour outcomes in mid-adolescence over ten years (2005 to 2015) using two key UK birth cohort studies. Importantly, the study investigated this range of outcomes within the same analytic framework, and employed methodological techniques to provide comparable estimates across the different health outcomes.

Prevalence of depressive symptoms, self-harm and parent reported mental health difficulties were all higher in 2015 compared to 2005, whereas anti-social behaviours were lower in 2015. Changes in these mental health outcomes were substantial, with a 6% increase (9% in 2005, 14.9% in 2015) in those above the threshold for depression and 20% decrease in those reporting physically assaulting anyone at age 14 (40.1% in 2005, ∼28% in 2015). Most antisocial behaviours reported were substantially lower in 2015 compared to 2005 and there was a sex interaction whereby the cohort difference was larger in females. Trends in externalising behaviours have been understudied in cohort comparisons and this data provides clear evidence for changes in antisocial behaviours in the decade between these cohorts.

The increase in internalising mental health problems was consistent by sex, suggesting that increases in psychological distress and self-harming behaviour are not increasing at higher rates in females (albeit higher base rates in females). This finding is in contrast to some older studies of adolescent trends that indicated that increases at the end of the 20^th^ century were more consistent and greater in females.^6,9^ For instance, previous research has reported odds in 2006 compared to 1986 at age 16 of 0.9 in males and 1.5 in females,^6^ compared to the increased odds in this study of ∼1.8 in both males and females in 2015 compared to 2005. It is striking that the rate of increase of high depressive symptoms is more than 60% in just one decade. Poor mental health at this age predicts a host of lifelong negative consequences such as poorer health, social and economic outcomes,^3,29^ and therefore this sharp increase should cause concern.

Results for health related behaviours were mixed with less young people having tried alcohol, binge drinking, smoking and having sex by mid adolescence in 2015 but being more likely to have later bedtimes and wake up earlier, to perceive themselves as overweight and to have higher BMIs. It is relevant to note that although fewer young people had tried smoking cigarettes in 2015, there was no cohort difference in the proportion smoking weekly at this age, although in absolute terms the number of individuals smoking weekly at age 14 was small (approximately 2% in both cohorts). In terms of sex differences in these cohort effects, the odds for some antisocial behaviours and ever trying alcohol in 2015 compared to 2005 were even lower in females compared to males, indicating that for some of these behaviours the decreasing prevalences over time were more marked in females.

The health-related behaviours identified in this study are all known risk factors for mental ill-health.^12-14,17^ In some instances the increasing trends in risky health behaviours such as decreasing sleep times, increasing weight, and perceived overweight status might help explain the increasing mental health difficulties experienced by adolescents. Where the trends are moving in opposite directions (substance use, antisocial behaviours), the interpretation becomes more complicated. It may suggest that the associations between these behaviours and mental health are not consistent over generations and might be changing over time. This is important with regards to trying to identify causal risk factors for poor adolescent health outcomes. Unexpected patterns such as those seen in our study, could indicate that associations between, for example, cannabis use and depression could be due to residual confounding rather than true causality. However, other factors not included in the study (e.g. family structure,^30^ parent mental health^31^) are also likely to have changed over the ten years of investigation, which may also impact on these associations. Understanding the dynamic relationships between health behaviours and mental health should be a priority as adolescent mental health problems increase, in order to identify suitable targets for interventions to prevent this upward trend from continuing.

In addition to effectively using two large contemporary birth cohort studies, the study makes several methodological advancements in improving our understanding of changing trends in UK adolescents. Variables in the two cohorts, where dissimilar, were carefully harmonised to ensure comparisons could be made. Unfortunately, this harmonisation could not be achieved for certain variables of interest (physical activity). Similarly, for other variables the harmonisation is imperfect either owing to different time periods of reference in the questions (e.g. self-harm) or availability only at a slightly different age in the ALSPAC cohort (e.g. sleep times) and this must be borne in mind when interpreting findings. In both these cases, however, the direction of bias is likely to be an underestimation of the increased poorer outcomes in 2015, for instance, with self-harm we estimate lifetime prevalence in ALSPAC and previous year prevalence in MCS. Although ALSPAC and MCS are large and detailed birth cohorts, one is a regional cohort (ALSPAC) and one is a national cohort (MCS), however, regional variation in these outcomes was estimated and was found to be minimal (<1% for mental health, sex, weight variables, <3% for substance use and sleep). Albeit employing multiple techniques to increase the comparability of the cohorts, it is possible that some of the differences observed are due to changes in demographic composition over the decade, differences in the study samples, or the different rates and predictors of attrition between the two studies. The nationally representative estimates for the MCS at age 14 indicate that across all the investigated variables the comparable estimates were slightly different from the nationally representative ones, highlighting the value of applying techniques to increase the comparability of these cohorts, but at the same time limiting the generalisability of our secular trend estimates to the UK as a whole.

There are a number of implications highlighted by our findings. Most importantly, the rapidly increasing prevalence of depressive symptoms, self-harm, parent-reported mental health problems, obesity and lesser sleep in adolescents over the past decade is an important finding, and the reasons why this has occurred need thorough investigation. Identifying further factors that have changed over the decade that might have resulted in UK young people having less support and being at higher risk should be undertaken as a public health priority. A further implication arising from our findings is that while certain mental health problems are increasing, other problems and health related behaviours, thought to predict poor mental health, are decreasing. Understanding the nature of these associations and their dynamic nature over time could be extremely valuable in identifying causal risk factors for mental health and potential targets for interventions. Identifying explanations for these high prevalences and changing time trends are key for preventing further increases in poor mental health and health outcomes for future generations of young people.

To conclude, in a large well-powered study across two key UK birth cohorts born a decade apart, depressive symptoms and self-harm behaviours have increased dramatically between 2005 and 2015. Adolescents are spending less time sleeping and have higher BMIs. In contrast, other health related behaviours such as substance use and antisocial behaviours have decreased over the same time period, suggesting that links between mental health problems and health related behaviours might be more complex and dynamic in nature than currently predicted. The data provide important evidence to understand health behaviours in millennials and how these are changing, permitting the planning of policy and public health provision.

## Acknowledgments

We are extremely grateful to all the families who took part in this study, the midwives for their help in recruiting them, and the whole ALSPAC team, which includes interviewers, computer and laboratory technicians, clerical workers, research scientists, volunteers, managers, receptionists and nurses.

The authors are grateful for the cooperation of the Millennium Cohort Study families who voluntarily participate in the study. They would also like to thank a large number of stakeholders from academic, policy-maker and funder communities and colleagues at the Centre for Longitudinal Studies involved in data collection and management.

## Funding

This work did not receive specific funding. The UK Medical Research Council and Wellcome (Grant ref: 102215/2/13/2) and the University of Bristol provide core support for ALSPAC. A comprehensive list of grants funding is available on the ALSPAC website (http://www.bristol.ac.uk/alspac/external/documents/grant-acknowledgements.pdf). The Millennium Cohort Study is supported by the Economic and Social Research Council and a consortium of UK government departments. The funders had no role in study design, data collection, data analysis, data interpretation, or writing of this report.

## Authors contributions

PP and SG, planned the study, analysed the data and prepared the manuscript for publication. Both PP and SG have full access to the data presented in this report act as guarantors for the paper.

## Declaration of Interests

All authors have completed the ICMJE uniform disclosure form at www.icmje.org/coi_disclosure.pdf and declare: no support from any organisation for the submitted work; no financial relationships with any organisations that might have an interest in the submitted work in the previous three years; no other relationships or activities that could appear to have influenced the submitted work.

## Supplementary Table S1

**Table S1.** Descriptive statistics for the key socio-demographic characteristics in the 2005 (ALSPAC) and 2015 (MCS) samples

|  | ALSPAC, N= 5627 | MCS full sample, N=11318 | MCS, N=5627<br>Propensity score matched | MCS, N= 11318<br>Entropy balanced |
| --- | --- | --- | --- | --- |
|  | Mean or % [95% CI] | Mean or % [95% CI] | Mean or % [95% CI] | Mean or % [95% CI] |
| Sex (female) | 50.72 [49.4,52.0] | 50.55 [49.6, 51.5] | 51.8 [49.5, 52.1] | 50.72 [48.2, 53.2] |
| Ethnicity (minority) | 3.87 [3.4,4.4] | 20.37 [19.6, 21.1] | 10.6 [9.8, 11.4] | 3.95 [3.6,4.4] |
| Age | 13.83 [13.83, 13.84] | 14.01 [14.25, 14.26] | 14.00 [13.99,14.01] | 13.83 [13.82, 13.85] |
| Maternal education (lower than GCSE/none/unclassified) | 20.19 [19.2, 21.3] | 16.2 [15.5, 16.9] | 17.03 [16.1,18.0] | 20.21 [18.1,22.5] |
| Maternal education (GCSE) | 35.37 [34.1, 36.6] | 35.6 [34.7, 36.4] | 38.38 [37.1,39.7] | 35.35 [33.1, 37.7] |
| Maternal education (A-levels) | 27.42 [26.3, 28.6] | 14.0 [13.3, 14.6] | 14.06 [13.2, 15.0] | 27.41 [24.9, 30.0] |
| Maternal education (degree or higher) | 17.03 [16.0,18.0] | 34.3 [33.4, 35.2] | 30.5 [29.3, 31.7] | 17.03 [15.7, 18.4] |
| Maternal age at birth | 29.66 [29.54, 29.78] | 28.97 [28.87, 29.08] | 29.33 [29.17, 29.49] | 29.66 [29.2, 30.1] |

